# Multiple myeloma immunoglobulin λ translocations portend poor prognosis

**DOI:** 10.1101/340877

**Authors:** Benjamin G. Barwick, Paola Neri, Nizar J. Bahlis, Ajay K. Nooka, Jonathan L. Kaufman, Vikas A. Gupta, Daniel Auclair, Jonathan J. Keats, Sagar Lonial, Paula M. Vertino, Lawrence H. Boise

**Affiliations:** Department of Hematology and Medical Oncology, Emory University School of Medicine, 1365 Clifton Rd. NE, Atlanta, GA 30322; Department of Radiation Oncology, Emory University School of Medicine, 1701 Uppergate Drive, Atlanta, GA 30322; Winship Cancer Institute, Emory University, 1365 Clifton Rd, Atlanta, GA 30322; Tom Baker Cancer Center, 3330 Hospital Drive, Calgary, AB, T2N 4N1; Multiple Myeloma Research Foundation, 383 Main Avenue, 5th Floor, Norwalk, CT 06851; Translational Genomics Research Institute, 445 North Fifth Street, Phoenix, AZ, 85004

## Abstract

Multiple myeloma is a malignancy of antibody-secreting plasma cells. Most patients benefit from current therapies, however, 20% of patients relapse or die within two years and are deemed ‘high-risk’. To better understand and identify high-risk myeloma, we analyzed the translocation landscape of 826 newly-diagnosed patients by whole genome sequencing as part of the CoMMpass study. Translocations at the IgL locus were present in 10% of myeloma patients, and corresponded with poor prognosis. Importantly, 70% of IgL translocations co-occurred with hyperdiploid disease, a marker of standard risk, which is routinely diagnosed clinically whereas IgL-translocations are not. Thus, it is likely that the majority of IgL-translocated myeloma is being misclassified. The IgL enhancer is among the strongest in myeloma cells, indicating it can robustly drive oncogene expression when translocated. Consistent with this, IgL-translocated patients failed to benefit from immunomodulatory imide drugs (IMiDs), which target the lymphocyte-specific transcription factor Ikaros. These data implicate the IgL enhancer as resistant to IMiD-inhibition, and when translocated, as a driver of poor prognosis.

## Introduction

Multiple myeloma is the second most common hematological cancer, which affects terminally differentiated antibody secreting B cells, known as plasma cells, and results in hypercalcemia, anemia, renal failure, and lytic bone lesions. Over the past decade, there have been significant Improvements in survival due to current therapies, which include autologous stem cell transplant ^1^, proteasome inhibitors ^2^, the immunomodulatory imide drugs (IMiDs), thalidomide ^3^, lenalidomide ^4,5^, and pomalidomide, and more recently monoclonal antibodies ^6,7^. Despite these advances, approximately 20% of patients relapse or die within two years of diagnosis ^8,9^. These patients are referred to as ‘high-risk’. Understanding how to identify and treat these patients as well as the mechanisms underlying the biology of high-risk myeloma is critical for improving outcomes.

Genetic analyses of myeloma over the last quarter century have revealed a bifurcation of founding genetic alterations with approximately half of myelomas containing an immunoglobulin heavy chain (IgH) translocation ^10^, which most commonly juxtapose the IgH enhancer with *CCND1* [t(11;14)], *WHSC1* [t(4;14); also known as *MMSET* and *NSD2*], *MAF* [t(14;16)], or *MYC* [t(8;14)]. The other half of myeloma harbor hyperdiploidy, which is an aneuploidy of chromosomes 3, 5, 7, 9, 11, 15, and 19 ^11^. Despite these seemingly simple explanations of the initiating events, the manifestation of myeloma at presentation is often confounded by a complex array of genetic alterations including amplification of chromosome 1q [amp(1q)], deletion of chromosome 13 [del(13)], deletion of chromosome 17p [del(17p)], dysregulation of MYC ^12^ and Cyclin D proteins ^13^, as well as mutations in common and disease-specific oncogenes (KRAS, NRAS, FAM46C, DIS3, BRAF, TRAF3, TP53) ^14^. Compounding the wide array of genetic abnormalities in myeloma is a daunting clonal heterogeneity, wherein selective pressures in the microenvironment and/or treatment promote the outgrowth of sub-clones harboring specific mutations that confer a survival advantage ^15^. Fortunately, modern combination therapies are mostly effective despite disease heterogeneity, with the majority of patients responding to frontline treatments that target plasma cell biology rather than specific genetic lesions ^16^.

Risk stratification currently uses the Revised International Staging System (R-ISS) ^17^, which incorporates the genetic abnormalities del(17p), t(4;14), and t(14;16) into the existing International Staging System (ISS) ^18^. While R-ISS improves prognostication, it fails to accurately identify all high-risk patients and newer therapies have mitigated the poor prognosis of some of the existing markers ^19^. Thus, reliable identification of high-risk patients remains a significant challenge.

To better understand the genetic basis of myeloma, and specifically high-risk disease, we investigated the genomic landscape of 826 newly diagnosed myeloma patients using long-insert whole-genome sequencing as part of the Clinical Outcomes in Multiple Myeloma to Personal Assessment (CoMMpass) study. These data identified recurrent translocations in 66.1% of newly diagnosed myeloma patients, with t(11;14), t(4;14), t(MYC), and other immunoglobulin translocations being the most common. While these translocations resulted in aberrant oncogene expression, very few were prognostic of altered outcome. The notable exception was patients with a translocation in the immunoglobulin light chain λ (IgL) locus, who experienced a significantly worse progression-free and overall survival, regardless of the juxtaposed locus or oncogene expression. The majority of patients with IgL translocations had hyperdiploid disease, which is a marker of standard risk, and thus could result in the misclassification of IgL-translocated [t(IgL)] patients. Furthermore, the poor prognosis of patients with t(IgL) was exacerbated by other high-risk markers such as amp(1q). The IgL locus is among the strongest enhancers in myeloma cells suggesting it may be resistant to therapeutic inhibition. Consistent with this t(IgL) patients did not benefit from IMiDs which target the lymphocyte-specific transcription factor IKZF1 ^20,21^, whereas other patients did. This suggests that the IgL enhancer is resistant to IMiD inhibition. These data identify IgL translocation as a multiple myeloma marker of poor prognosis independent of other genetic abnormalities with implications for the diagnosis and treatment of myeloma.

## Results

### The translocation architecture of newly diagnosed multiple myeloma

A comprehensive analysis of structural variants in multiple myeloma was conducted using long-insert whole-genome paired-end sequencing performed on DNA isolated from CD138+ myeloma cells to determine cancer-specific somatic mutations, and normal peripheral blood to determine the germline genome as part of the CoMMpass study (NCT01454297). CoMMpass is a longitudinal study of 1,000 newly diagnosed myeloma patients enrolled from North American and European collection sites. In total, samples from 826 newly diagnosed patients were subjected to long-insert sequencing with an average of 304 million paired-end reads per specimen (8.3x coverage), and an average fragment size of 846 bp, thus spanning the genome with 40.9x coverage. Structural variants including deletions, duplications, inversions, and translocations were identified using DELLY ^22^ (**Table S1**). This analysis identified a median of 21 structural variants and 3 translocations in newly diagnosed myeloma (Fig. 1a). Deletions, duplications, and translocations corresponded with worse progression-free (PFS) and overall survival (OS), with translocations being the most significant (Fig. 1b). These data are consistent with the concept of genomic instability being associated with higher risk ^9^.

**Figure 1.**
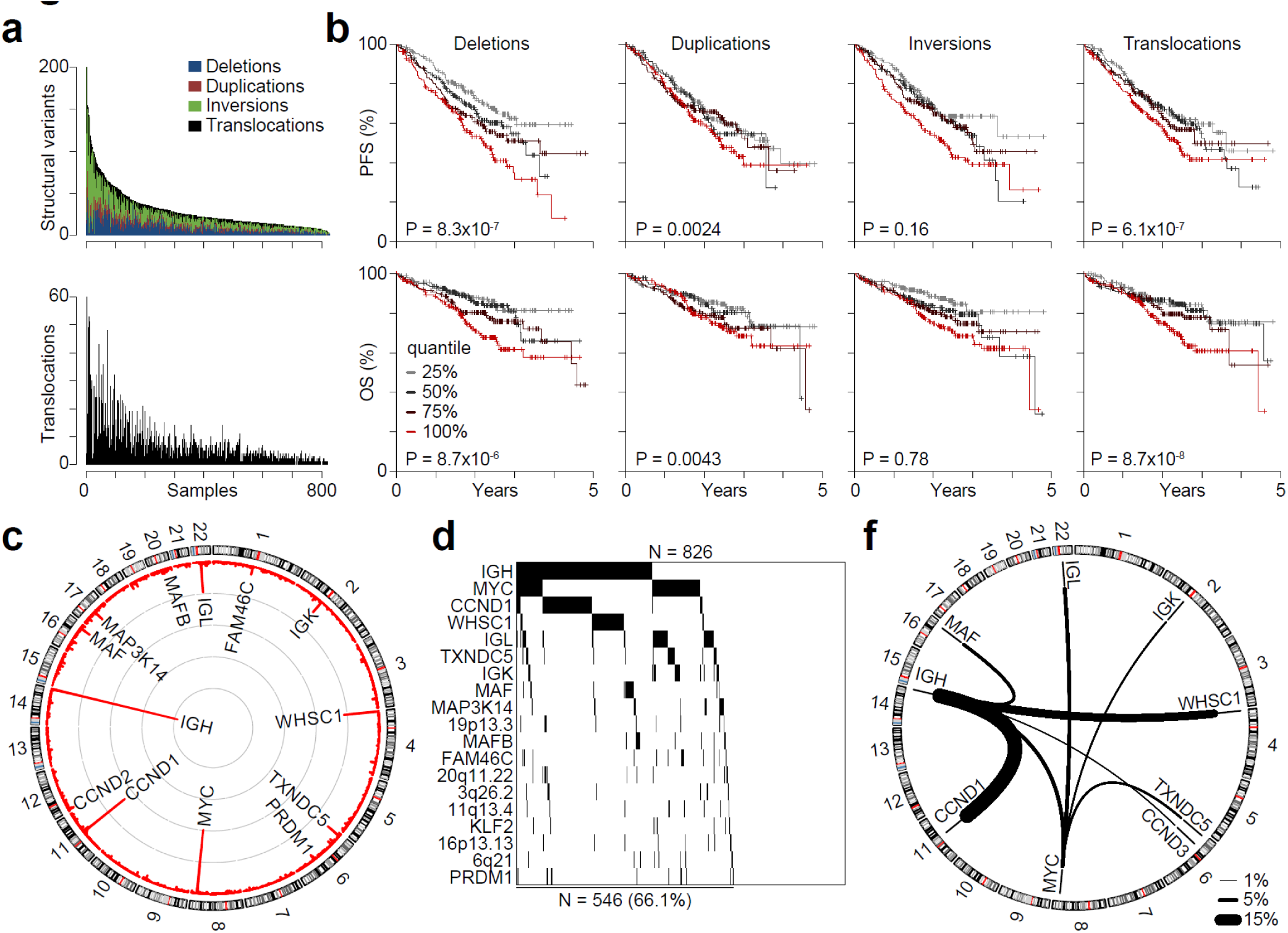
Structural variants correspond with poor prognosis. **a,** The number of all somatic structural variants (top) and translocations (bottom) per sample in 826 newly diagnosed myeloma specimens ordered by the total number of structural variants. The structural variant plot is cropped at 200 structural variants with 4 samples having more than 200. **b,** Progression-free (PFS) and overall survival (OS) for patients stratified into quartiles based on the number of deletions (left), duplications (middle left), inversions (middle right), and translocations (right). **c,** Circos plot of translocation frequency in 826 newly diagnosed myeloma patients. Chromosomes ideograms are represented and labelled on the outside and the frequency of translocations are denoted in red on the inside with gray concentric circles denoting 10 percentiles per megabase. **d,** Waterfall plot of translocated regions showing regions translocated in ≥2% of newly diagnosed patients. **e,** Circos plot of translocated regions prevalent in ≥1% of the newly diagnosed population. Thickness of lines connecting two regions denotes the frequency of the translocation (see key bottom left). P-values were determined using a Cox proportional hazards regression Wald test as a function of the number of structural events.

Commonly translocated regions were identified using a 1 Mb window incremented by 0.5 Mb across the genome. Frequent translocations included regions proximal to *IgH* (41.4%), *MYC* (22.5%), *CCND1* (17.3%), *WHSC1* (11.0%), *IgL* (9.8%), *TXNDC5* (4.9%), *IgK* (4.3%), and *MAF* (4.1%) (Fig. 1c). In total, 19 regions were translocated at a frequency of 2% or more with 66.1% of newly diagnosed patients having at least one of these rearrangements (Fig. 1d). Common reciprocal translocations were determined by comparing the frequency of translocations within a 1 Mb region, which revealed 8 translocation types present in at least 1% of newly diagnosed patients (Fig. 1e). As expected, the most common included the IgH translocations t(11;14) (16.1%), t(4;14) (10.4%), t(8;14) (3.7%), and t(14;16) (3.4%) as well as MYC translocations to the IgL (4.0%) and immunoglobulin κ (IgK) (1.9%) loci. These data define the repertoire of translocations in newly-diagnosed myeloma and resolve the breakpoints to within 1 kb.

### The breakpoint of IgH translocations indicate multiple mechanisms

The pathologic significance of translocations was studied starting with the most frequent region, the IgH locus. The most common IgH translocations included t(11;14), t(4;14), t(8;14), and t(14;16), which accounted for 283 of 342 (82.7%) IgH-translocated patients (Fig. 2a). These translocations resulted in aberrant upregulation of the transposed gene (**Fig. S1a**). IgH translocations preferentially occurred near the class switch recombination (CSR) regions located at the 5’ edge of the constant heavy chains μ (M), γ_1_ (G_1_), α_1_ (A_1_), γ_2_ (G_2_), α_2_ (A_2_), and γ_3_ (G_3_) (Fig. 2b; top). However, there were distinct differences between the IgH translocation species. For instance, t(11;14), t(4;14), and t(14;16) translocations were primarily located at CSR regions, with t(11;14) translocations more likely to occur at the G_1_, A_1_, G_2_, A_2_, G_3_ CSR regions, and t(4;14) almost exclusively at the M switch region. In contrast, t(8;14) and other IgH translocations preferentially occurred at extragenic regions (Fig. 2b). To determine if there were differences in breakpoint location between the IgH translocation species, translocations arising from the CSR, VD, and extragenic regions were quantified and biases in location were expressed as an odds ratio of overlap for each region and translocation (**Fig. S1b**). These data further indicated that t(11;14), t(4;14), and t(14;16) translocations preferentially occurred at the CSR regions, with t(4;14) showing the strongest bias. Conversely, t(8;14) and other IgH translocations preferentially occurred at extragenic regions (**Fig. S1b**, right). These data suggest that the majority of IgH translocations occurred during germinal center B cell CSR, consistent with previous observations ^10^, except for IgH-MYC translocations, which may occur by a different mechanism. Nonetheless, despite the distinct etiologies of the aforementioned translocations, patients harboring these abnormalities experienced outcomes commensurate with each other and other myeloma patients (data not shown).

**Figure 2.**
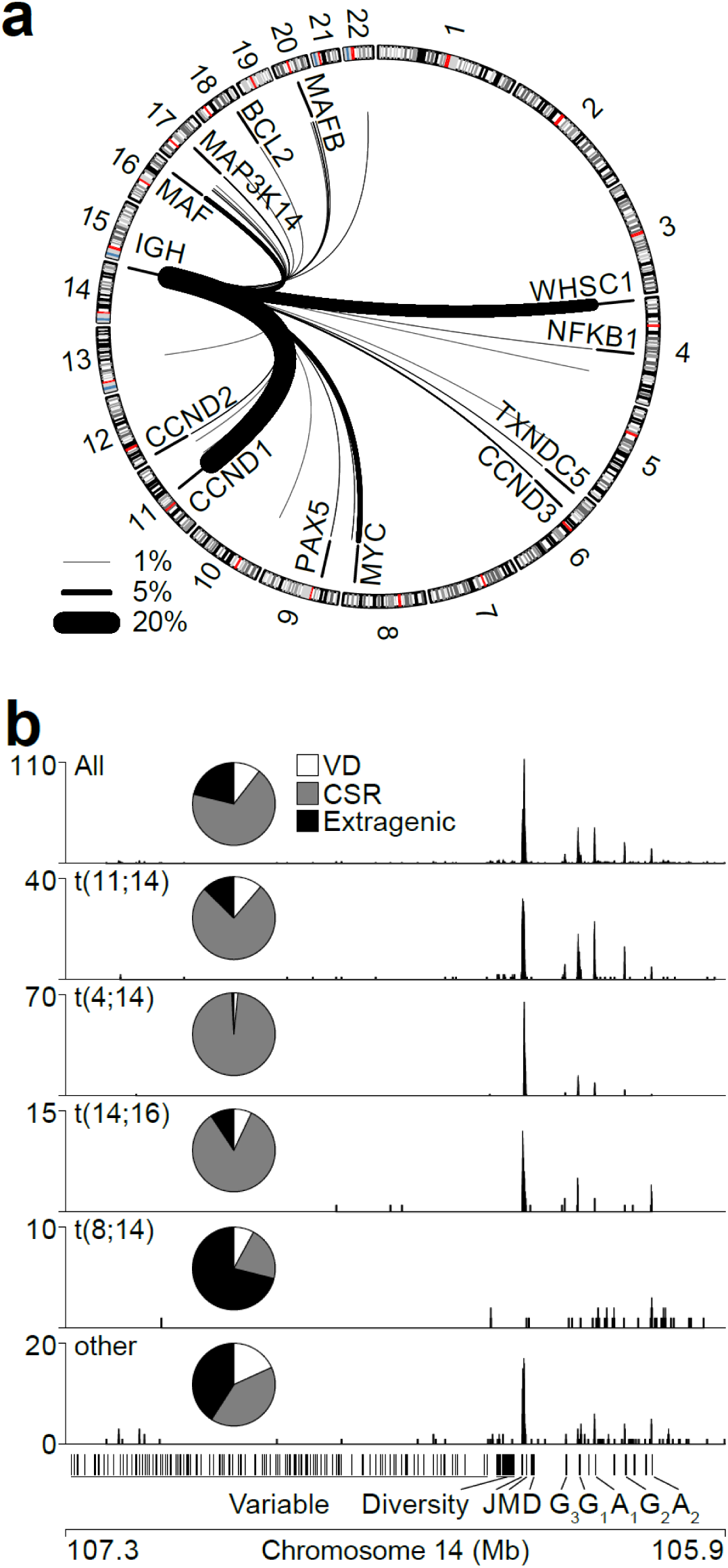
IgH translocations have distinct etiologies. **a,** Circos plot of IgH translocations in newly diagnosed myeloma patients (N=826). The frequency of each type of translocation is denoted by the thickness of the line (key bottom left). **b,** Genome plots of translocations across the IgH region for all IgH translocations (top) or specific types of IgH translocations (below). Inset are pie charts depicting the percentage of translocations that occur in the variable and diversity (VD), class switch recombination (CSR) (+/− 2.5kb), and extragenic regions.

### MYC translocations are juxtaposed throughout the genome

Since IgH-MYC translocations had a pattern of breakpoints that was distinct from most other IgH translocations, the MYC translocation landscape was determined for newly diagnosed myeloma. MYC translocations occurred in 186 (22.5%) of 826 patients, and was juxtaposed to a large number of regions (Fig. 3a). The most common regions included IgL [t(8;22); N=33], IgH [t(8;14); N=31], TXNDC5 (N=19), IgK [t(8;2); N=16], FOXO3 (N=8), and FAM46C (N=6). Unlike IgH, which was translocated primarily to a few specific loci, a larger proportion of MYC translocations (N=73; 39.2%) occurred throughout the genome. Also, unlike IgH translocations, MYC translocation breakpoints were clustered across two broad regions, one centered on MYC and the other 600 kb to the telomeric side of *MYC* (Fig. 3b). MYC translocations occurred near regions enriched for H3K27ac, a histone modification present at active enhancers, in the myeloma cell line MM.1S (Fig. 3b). Analysis of copy-number alterations (CNA) across the MYC locus identified focal amplifications that commonly occurred near the most frequently translocated regions, which were more often observed among myelomas that contained a MYC translocation (Fig. 3b). Indeed, analysis of MYC CNV data derived from exome sequencing in 754 samples, confirmed that MYC translocations corresponded with copy number gains in the region, and showed that among the 118 (15.6%) myelomas containing MYC amplification, 50% also had a MYC translocation (Fig. 3c). This is in contrast to non-MYC amplified myeloma which only contained MYC translocations in 17.8% of cases (Fig. 3c, see inset table). Both genomic alterations resulted in increased MYC expression, and there was no difference in expression between MYC amplified myelomas and those with different types of MYC translocation (Fig. 3d). Nor was there a difference in expression between MYC translocations that occurred proximal or distal to MYC (data not shown). Interestingly, despite similar levels of MYC expression, distinct MYC-translocation partners exhibited differences in outcome. Indeed, among MYC translocations, MYC-IgL t(8;22) translocations had the worst PFS and OS (Fig. 3e). These data indicate that aberrant MYC expression resulting from MYC amplification or translocation is a common feature of myeloma, but the MYC-IgL translocated subset is unique among MYC alterations in that it portends a poor prognosis.

**Figure 3.**
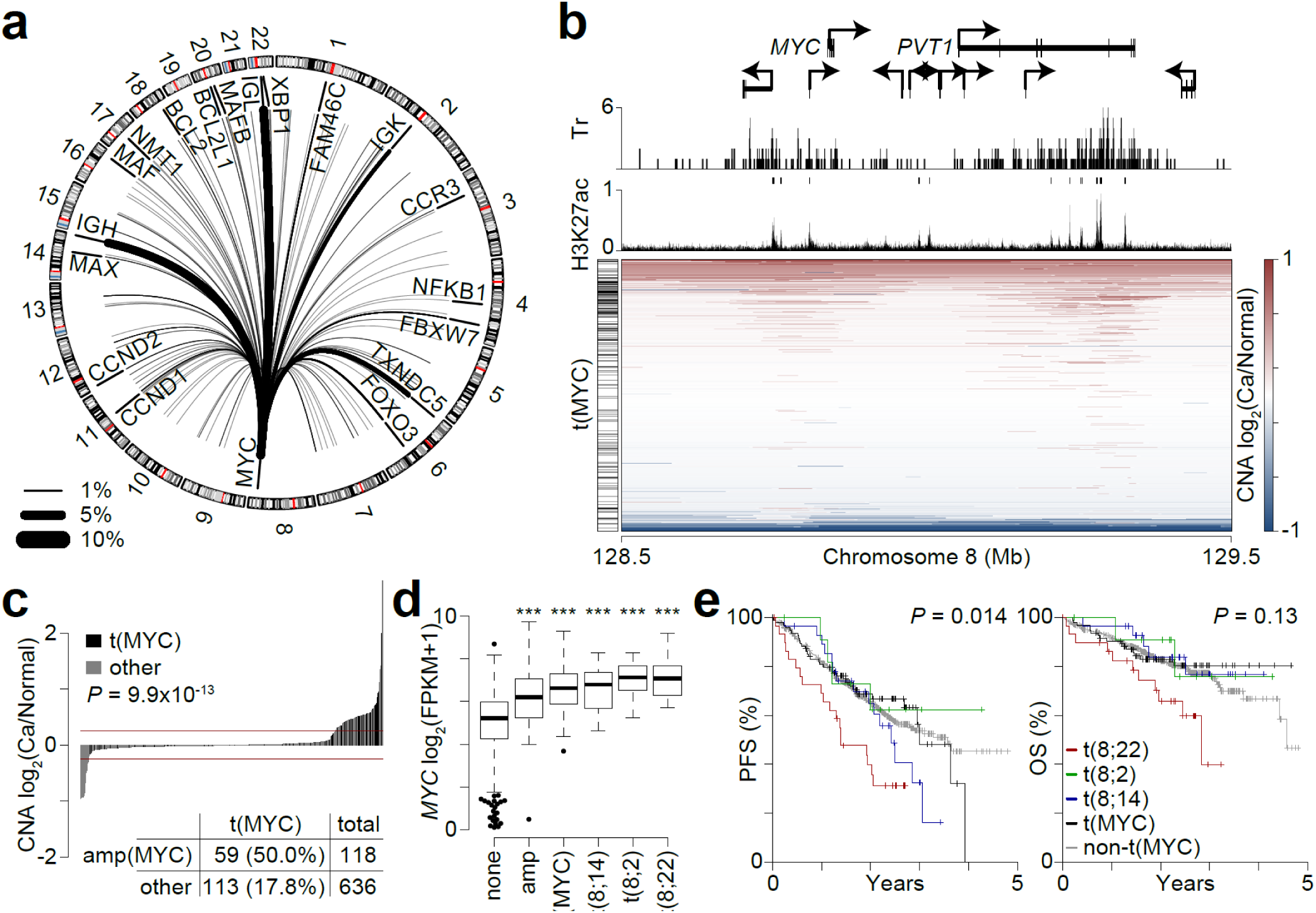
MYC translocations correspond with aberrant expression and copy number gains. **a,** Circos plot of MYC translocations in newly diagnosed myeloma (N=826) where the frequency of each translocation is denoted by the thickness of the line (key bottom left). **b,** Genomic plot of translocations at the MYC locus. Select genes are labeled (top), with translocations (Tr; top middle), and H3K27ac ChIP-seq data in MM.1S (bottom middle) shown. H3K27ac-enriched regions are denoted immediately above the H3K27ac track. Copy number alterations (CNA) are shown (below) as a log2 ratio of myeloma to normal (Ca/Normal) and are sorted by average CNA across the 1MB region shown. **c,** CNAs at the MYC locus for newly diagnosed myeloma patients with translocation and exome copy number data (N=754). Myelomas with a MYC translocation are denoted in black and the number of myelomas with translocated and amplified MYC are shown in the table inset. Red lines denote the CNA thresholds (+/−0.2 log_2_). **d,** Gene expression of MYC in newly diagnosed myelomas with translocation, CNA, and gene expression data (N=595) for patients with no MYC translocations or amplifications (none; N=406), MYC amplification but no translocation (amp; N=48), MYC translocation to a non-immunoglobulin gene (t(MYC); N=79), MYC-IgH translocation (t(8;14); N=24), MYC-IgK translocation (t(8;2); N=12), and MYC-IgL translocation (t(8;22); N=26). *\*\*\*P* < 0.001, analysis of variance with Tukey’s post-hoc test. Boxplots show the median and quartiles with the whiskers extending to the most extreme data point within 1.5 times the interquartile range. **e,** Progression-free (PFS; left) and overall survival (OS; right) for patients stratified by MYC translocation type: non-t(MYC) (N=640), MYC-other (N=106), t(8;14) (N=31), t(8;2) (N=16), and t(8;22) (N=33). P-values were calculated using a Cox proportional hazards Wald test and denote significant differences in survival between the categories shown.

### Translocation of IgL is an independent marker of poor prognosis

The above findings prompted us to examine IgL translocations in more detail. IgL was translocated in 9.8% (N=81/826) of newly diagnosed myeloma with 40.7% of IgL translocations being juxtaposed to MYC and the remaining were scattered throughout the genome, but included recurrent translocations proximal to *MAP3K14*, *CD40*, *MAFB*, *TXNDC5*, *CCND1*, *CCND2*, and *CCND3* (Fig. 4a). As suggested by the MYC analysis, t(IgL) patients experienced a worse PFS and OS as compared to non-t(IgL) myeloma (Fig. 4b). This was further confirmed through permutation analysis and was not attributable to differences in patient population as t(IgL) patients were of similar age, disease stage, sex, race, had similar levels of serum M-protein and β2-Microglobulin, and were treated with similar therapies as compared to patients without t(IgL) (**Fig. S2a-g**). Multivariate survival analysis was performed to identify any potentially confounding factors, and t(IgL) remained a significant marker of poor outcome when considering other known prognostic variables including age and stage (ISS) (**Fig. S2h**). Additionally, survival of patient’s with t(IgL) was compared to patients with other immunoglobulin translocations (**Fig. S3a**), which indicated that patient’s with t(IgL) had a worse outcome than those with t(IgH) or t(IgK). Finally, patients with IgL-MYC translocations and IgL-non-MYC translocation were compared to MYC-non-IgL translocation and non-IgL-non-MYC translocations (**Fig. S3b**), which showed that patients with IgL translocations had a worse prognosis, regardless of the partner. Taken together, these data identify t(IgL) as an independent marker of poor prognosis regardless of the transposed loci.

**Figure 4.**
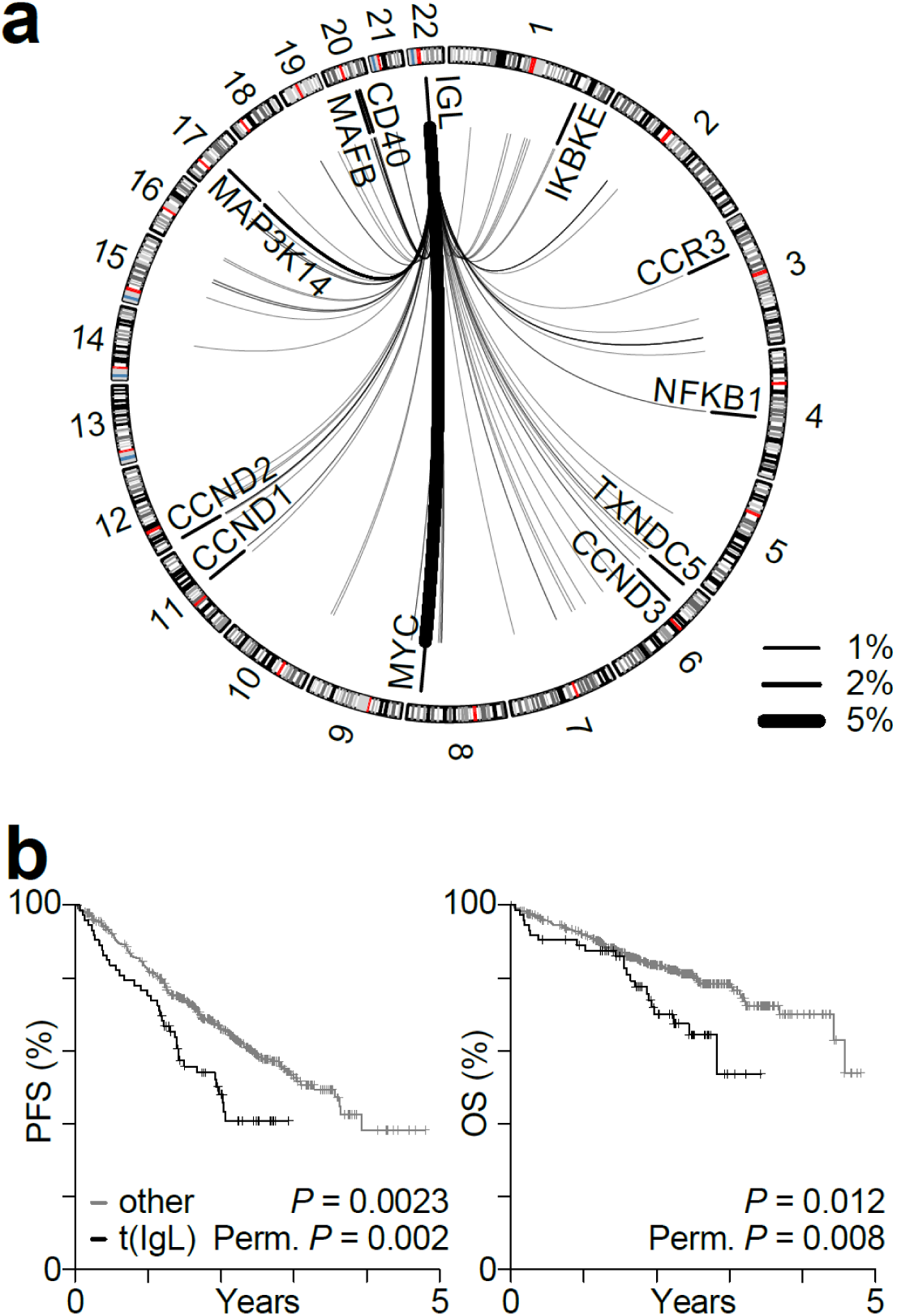
IgL translocations portend poor prognosis. **a,** Circos plot showing the repertoire of IgL translocations in newly diagnosed myeloma where line thickness denotes frequency (key bottom left). **b,** Kaplan-Meier analysis of IgL translocated [t(IgL)] patients (N=81) as compared to non-t(IgL) (N=745) for progression-free (PFS; top) and overall survival (OS; bottom). P-values were calculated using a Cox proportional hazards Wald’s test or permutation based P-value with 1,000 permutations based on the hazard ratio.

To determine whether there might be other molecular features that contribute to the poor outcomes of t(IgL) patients, the mutational repertoire of t(IgL) myeloma was interrogated using high-depth exome sequencing on 814 of the 826 specimens characterized by long-insert whole genome sequencing. This analysis indicated that there was no difference in the total and nonsynonymous mutational burden among t(IgL) myeloma compared to those with IgK and IgH translocations or no immunoglobulin translocation (**Fig. S4a,b**). The frequency of specific mutations types also indicated that t(IgL) myeloma had a similar mutational spectrum as other myelomas where all myelomas had a higher rate of transitions (C>T and A>G) than transversions (**Fig. S4c**). Analysis of specific mutations indicated that KRAS and NRAS were the most frequently mutated genes, as previously reported ^14^ (**Fig. S4d**). Although t(IgL) myeloma had a slightly higher rate of mutations in RYR1, CSMD1, USH2A, and FLG than other myelomas, none of these were statistically significant, and in general, t(IgL) myeloma contained a similar frequency of mutations in the most commonly mutated genes as compared to all other myelomas. Most mutations had a marginal impact on prognosis as assessed by a univariate analysis (data not shown). Furthermore, a bivariate analysis considering each mutation in combination with t(IgL), identified t(IgL) as prognostic of poor outcome independent of any other common mutation (**Fig. S4e**, right).

### IgL-translocated myeloma is not defined by gene expression subtype

To gain insight into the pathogenesis of IgL translocation, the relationship between t(IgL) and gene expression subtypes were determined using consensus clustering ^23^ on 654 samples for which whole genome sequencing and RNA-seq data were available. Similar to previous reports ^24^ this identified 7 gene expression subtypes, where samples within a given subtype were highly correlated with each other but not with samples from other subtypes (Fig. 5a). Both up- and down-regulated genes in each cluster were determined (**Fig. S5a; Table S2**), and annotated using gene set enrichment analysis (GSEA; **Table S3**) ^25^. This matched 5 of the 7 expression subtypes identified here with the MMSET (MS), hyperdiploid (HY), proliferation (PR), Cyclin D (CD), and MAF (MF) subtypes previously defined ^24^ (**Fig. S5b**). Group six corresponded with genes dysregulated in a MYC and BCL2L1 driven mouse model of multiple myeloma ^26^ as well as expression of more innate B cell and/or myeloid genes and is subsequently referred to as the myeloid subtype (MY). Finally, the seventh group modestly corresponded to the low bone (LB) disease signature, which was previously noted to have non-distinct gene expression pattern ^24^ (**Fig. S5b**; Fig. 5a). As expected, gene expression groups corresponded with genetic abnormalities where the MS subtype harbored t(4;14), the HY gene expression subtype corresponded with genetic hyperdiploidy, the PR subtype contained amp(1q), the CD subtype had t(11;14), and MF had t(14;16) (**Fig. S5c**, Fig. 5a - see annotation above). Notably, IgL translocations were found in every expression subtype with a modest but non-significant enrichment in the HY and PR gene expression subtypes and depletion in the CD subtype (Fig. 5b).

**Fig 5.**
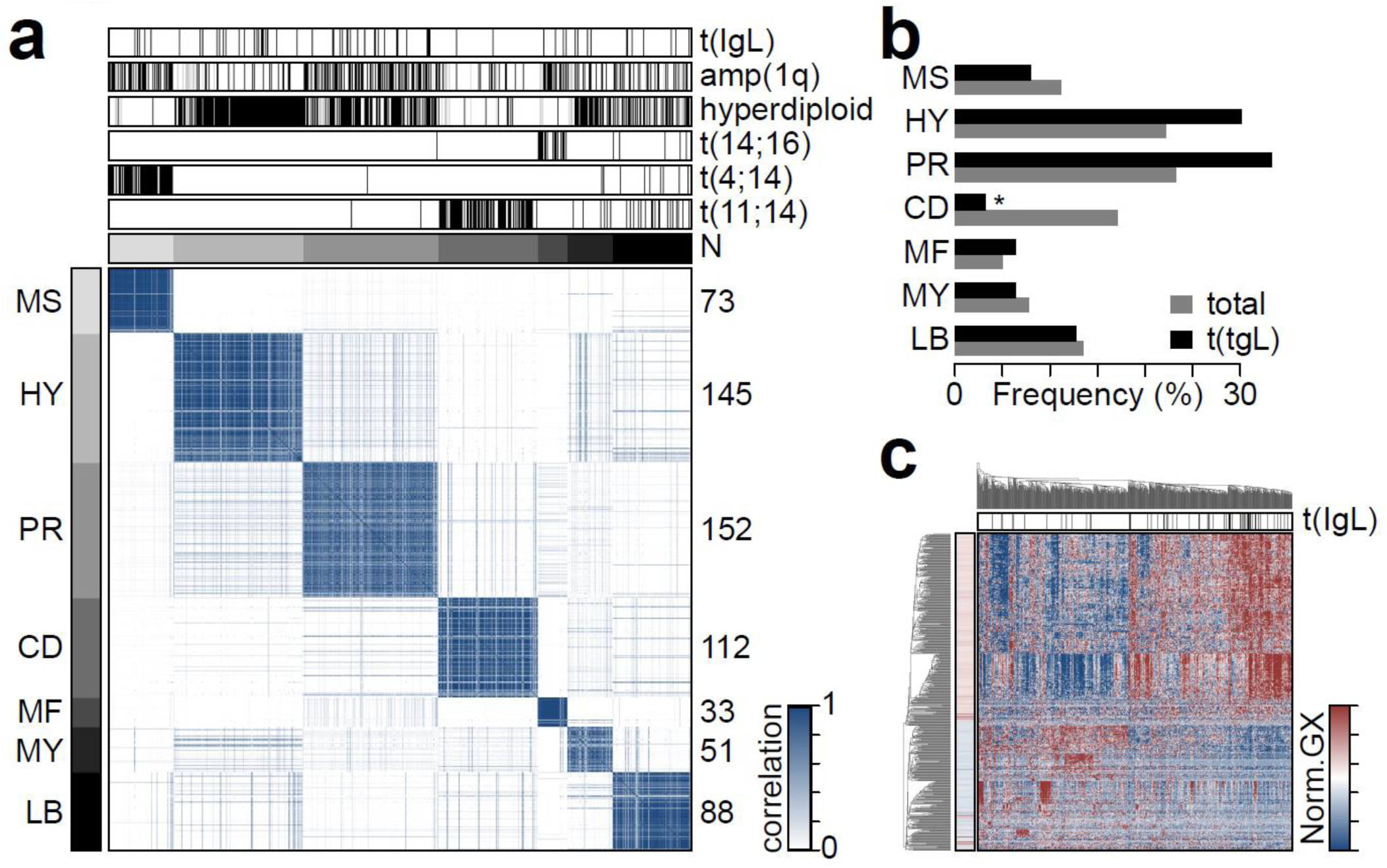
IgL translocation occur in all gene expression subtypes. **a,** Consensus clustering of 654 newly diagnosed myelomas with whole genome sequencing and RNA-seq. Expression subtypes are denoted by gray annotation bars (top and left), and labels correspond with those defined by Zhan et al. ^24^. The number of samples are denoted on the right. **b,** Frequency of gene expression subtypes in all myelomas (gray) and in t(IgL) myeloma (black). **c,** Heatmap of genes differentially expressed in t(IgL) myeloma as compared to others. Samples (columns) are clustered with t(IgL) status annotated (top; black) and gene significance denoted (left). *P <0.05; Fisher’s exact test.

To identify putative molecular mechanisms contributing to t(IgL) pathogenesis, genes differentially expressed in t(IgL) were determined, revealing 330 upregulated and 186 downregulated genes in t(IgL) myeloma as compared to non-t(IgL) myeloma (**Table S4**; FDR <0.01). Expression of these genes modestly aggregated t(IgL) myelomas using hierarchical clustering (Fig. 5c), but clearly grouped many non-t(IgL) with t(IgL) myelomas, suggesting that t(IgL) myeloma is not clearly defined by a baseline gene expression signature. GSEA identified several gene sets and pathways corresponding with t(IgL) differentially expressed genes including overexpression of ribosomal genes *RPS2* and *RPL4* that are involved in protein translation, components of energy metabolism including oxidative phosphorylation and respiratory electron transport chain, such as the NADH ubiquinone oxidoreductase complex component *NDUFAF4,* and genes upregulated downstream of MYC, including MYC itself, and the eukaryotic translation elongation factor *EIF3J* (**Fig. S6a**, top; **Table S5**). These differentially expressed genes, which while statistically significant, showed only modest differences between t(IgL) from non-t(IgL) myeloma (**Fig. S6b**, top). Likewise, the genes most downregulated in t(IgL) myeloma which included those normally repressed during B cell to plasma cell differentiation as well as genes involved in cytokine and chemokine signaling (**Fig. S6a**, bottom), showed only subtle differences between t(IgL) and non-t(IgL) myeloma (**Fig. S6b**, bottom). These data indicate that t(IgL) occurs across all gene expression subtypes of myeloma and indicate that translocation of IgL does not drive a unique gene expression program.

### IgL translocations co-occur with hyperdiploid disease

As t(IgL) was not associated with any specific mutations and had only modest correlations with gene expression, we investigated structural variants that might be associated with t(IgL). The frequency of specific loci translocated directly to IgL as well as those that co-occur with t(IgL) were compared to myeloma with IgH and IgK translocations (**Fig. S7a**). This indicated that IgH was more frequently translocated to *CCND1, WHSC1,* and *MAF,* whereas IgK and IgL were more frequently translocated to MYC. Approximately 40% of both IgK and IgL translocations were to MYC and another 20% of both t(IgK) and t(IgL) myelomas contained a MYC translocation but to a different locus (**Fig. S7b**). t(IgL) myeloma contained very few unique translocation partners, in that most loci transposed to *IgL* were also transposed to *IgK* or *IgH.* The most common translocations unique to *IgL* were MAP3K14 and 3q26.2, which only accounted for 7.4% and 4.9% of t(IgL) myeloma, respectively. Thus, these data suggest that the unique pathologic effects of IgL translocation are directly related to the IgL locus and not necessarily to a gene dysregulated by IgL transposition.

Other structural variants were also interrogated including deletions, duplications, and inversions, and showed that t(IgL) myeloma had a slightly larger number of duplications than t(IgH) myeloma (**Fig. S7c**). Thus, common CNAs were annotated for 754 newly diagnosed myelomas for which exome sequencing and whole genome sequencing data were available. A heatmap of CNAs identified a bifurcation in samples, where approximately half (N=382; 50.7%) showed a characteristic hyperdiploid pattern, exhibiting aneuploidy of odd numbered chromosomes including 3, 5, 7, 9, 11, 15, 17, and 19 (Fig. 6a). Hyperdiploid myelomas were mostly mutually exclusive with IgH translocations t(11;14), t(4;14), and t(14;16) as previously reported ^27^. t(IgL) was found in both groups but was more common in hyperdiploid disease (Fig. 6a). Indeed, del(17p), del(1p), amp(1q), del(13q) were found at similar frequencies in t(IgL) as in all other myelomas, but hyperdiploid disease was found in 70.4% of t(IgL) myeloma, which was a significant overrepresentation (Fig. 6b). Multivariate survival analysis of t(IgL) and the aforementioned CNAs indicated that t(IgL) was prognostic of poor PFS and OS even when accounting for all the other common CNAs, some of which have themselves been independently associated with a poor prognosis ^28^ (Fig. 6c). Notably, patients with hyperdiploid disease generally experience better outcomes ^28^, but this was not the case for patients with t(IgL) and hyperdiploid disease, who had significantly worse outcomes, as compared to other patients with hyperdiploid disease or other myeloma (Fig. 6d). These data identify IgL translocation as a poor prognostic prevalent in 14.9% of hyperdiploid myeloma that would otherwise be classified as standard-risk. Thus, a large portion of t(IgL) myeloma is likely being misclassified.

**Fig 6.**
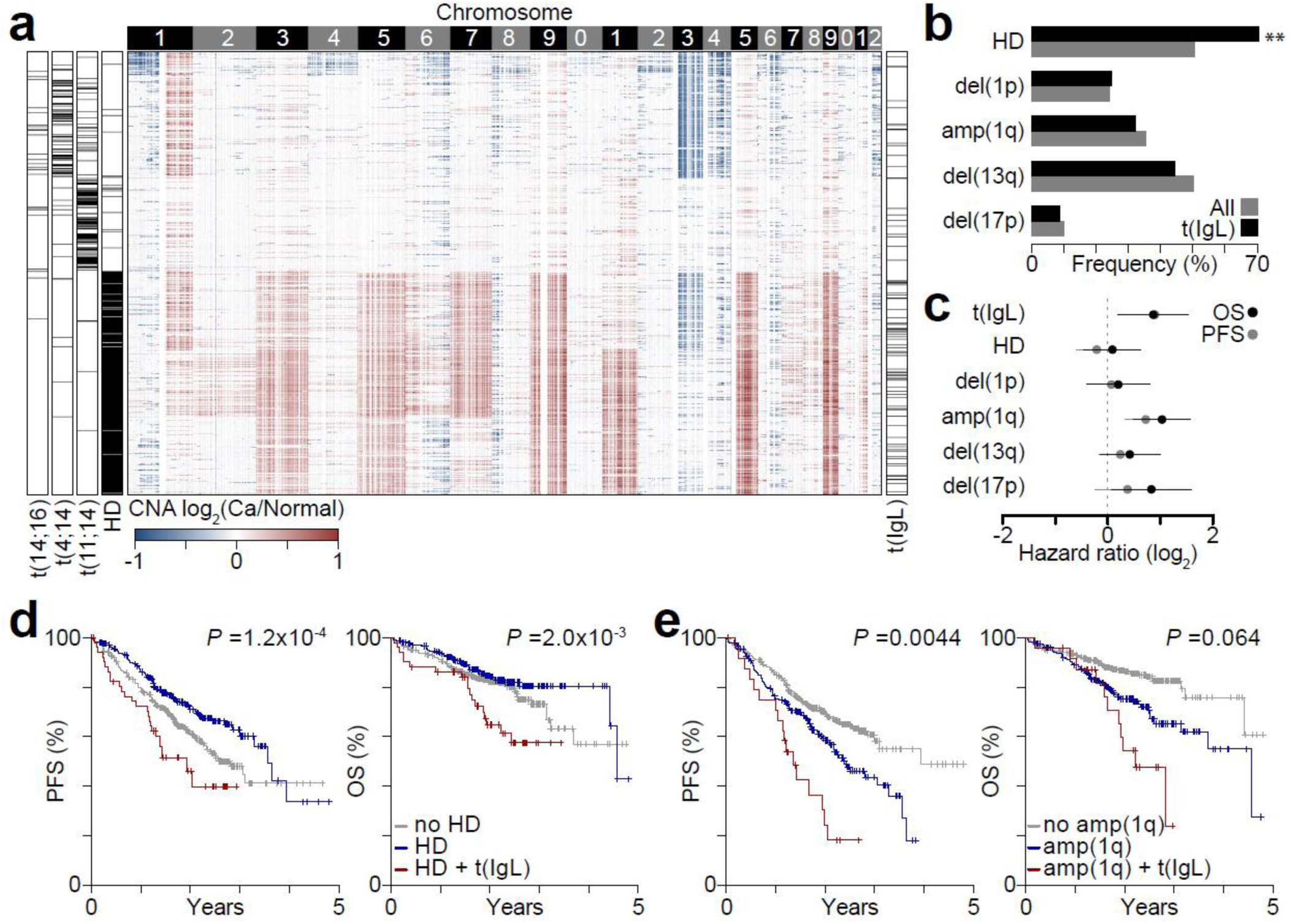
IgL translocation co-occurs with hyperdiploid disease. **a,** Heatmap of copy number alterations (CNA) in 754 newly diagnosed myelomas (rows) with genomic location (columns; 100 kb bins) labelled by chromosome (top). Hyperdiploid (HD) disease, and t(11;14), t(4;14), and t(14;16) translocations are annotated (left) as well as t(IgL) (right). **b,** Frequency of common CNAs in all newly diagnosed myeloma (gray) and myelomas with t(IgL) (black). *\*\*P* <0.01, Fisher’s exact test. **c,** Progression-free (PFS; gray) and overall survival (OS; black) hazard ratios determined by multivariate survival analysis of common CNAs and t(IgL). 95% confidence intervals are shown. **d,** Kaplan-Meier survival analysis of patients without HD (gray), with HD (blue), or with HD and t(IgL) (red). **e,** Same as part **d** except shown for amplification of 1q [amp(1q)]. *P*-values denote the survival difference between the specified CNA (blue) and the CNA + t(IgL) (red) determined by a Cox proportional hazards Wald’s test.

The above multivariate analysis also indicated amp(1q) as an independent marker of poor prognosis. Indeed, amp(1q) was associated with worse PFS and OS as compared to non-amp(1q) myeloma (Fig. 6f), similar to previous reports ^28^. Interestingly, the poor prognosis associated with amp(1q) was further exacerbated by the concurrent presence of t(IgL), which resulted in a median PFS and OS of 1.37 and 2.23 years, respectively (Fig. 6f). This suggests that the pathogenic effects of t(IgL) and amp(1q) have additive effects on myeloma outcomes.

### The IgL locus is a super-enhancer

The above data indicate that t(IgL) is an independent marker of poor prognosis regardless of translocation partner or other known molecular features, which suggests a factor intrinsic to the IgL locus may be mediating this pathology. One possibility is that the IgL enhancer is particularly potent in driving oncogene expression. Thus, we analyzed the chromatin structure of the IgL locus using ChIP-seq data in the myeloma cell line MM.1S ^29^. These data indicated that the IgL locus contained several regions with the activating histone modification histone 3 lysine 27 acetylation (H3K27ac), and these coincided with MED1, BRD4, and MYC occupancy (**Fig. S8a**). Although a similar 200 kb region at the 3’ end of the IgH locus also contained these marks, the IgL enhancer appeared to be more densely clustered than that of the IgH (**Fig. S8b**). Analysis of H3K27ac- or MED1-defined ‘super enhancers’ indicated that multiple enhancers at the IgL locus were among the biggest in the genome (**Fig. S8c**). Indeed IgL super-enhancers were larger than those of the IgH locus. These data indicate that the IgL locus contains several super-enhancers that when translocated likely serve as potent inducers of juxtaposed oncogenes.

### t(IgL) patients do not benefit from treatment with IMiDs

Given that t(IgL) patients experienced worse outcomes than other patients we hypothesized that this may be due to certain therapies being ineffective against t(IgL) myeloma. Treatment of myeloma in the CoMMpass trial was determined by physician’s choice, resulting in a diversity of front line treatments including proteasome inhibitors (79.3% of patients), dexamethasone (79.1%), IMiDs (62.2%), melphalan (33.5%), and cyclophosphamide (30.1%) (**Fig. S2**). We tested the outcome of t(IgL) patients in the context of these agents and this indicated that t(IgL) patients showed a similar PFS and OS regardless of IMiD treatment (Fig. 7a, compare blue and red lines). This is in stark contrast to non-t(IgL) patients who derive clear benefit from treatment with IMiDs (Fig. 7a, compare gray and green lines). Indeed, a significant survival benefit was provided by IMiDs for non-t(IgL) patients whereas no IMiD survival benefit was realized for patients with t(IgL) who had a PFS and OS similar to patients who did not receive an IMiD (Fig. 7a, see table right). In contrast, patients with IgH translocations who received IMiDs did significantly better than t(IgH) patients that did not (Fig. 7a). Importantly, bootstrap sampling of t(IgH) patients based on the number of t(IgL) patients indicated that t(IgH) patients who received IMiDs had a better progression-free survival than t(IgL) patients (PFS *P*=0.045; OS *P*=0.163). Thus, the lack of IMiD benefit observed in t(IgL) patients is not simply due to the smaller number of t(IgL) patients. Together, these data indicate that IMiDs are less affective against t(IgL) myeloma than other myelomas.

**Fig 7.**
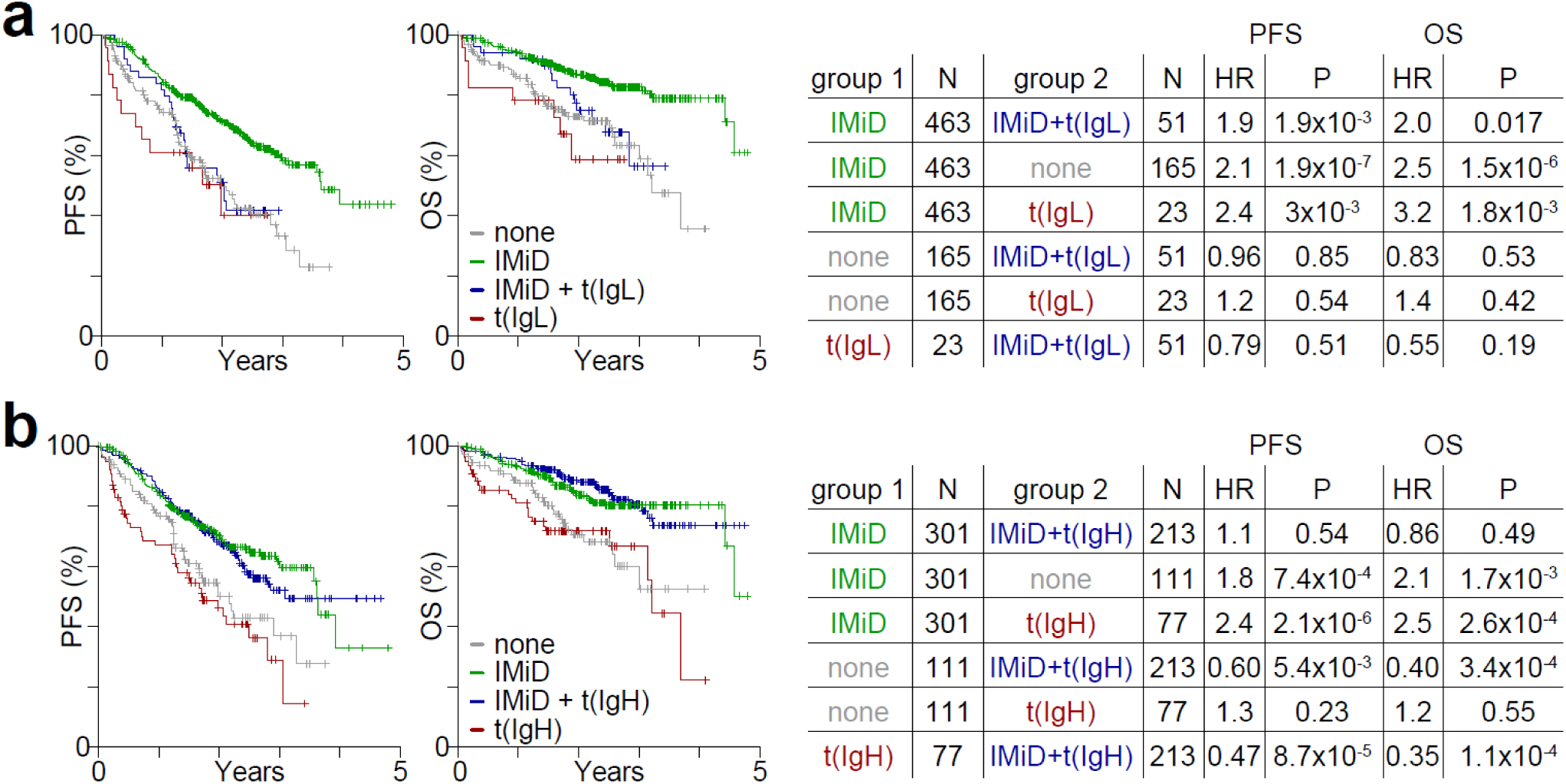
IgL translocated myelomas do not benefit from IMiDs. **a,** Progression-free (PFS; left) and overall survival (OS; middle) Kaplan-Meier survival curves for patients stratified by IMiD therapy and t(IgL) status: none (gray; no IMiD and no t(IgL)), IMiD (green; IMiD and no t(IgL)), IMiD+t(IgL) (blue), and t(IgL) (red; no IMiD and t(IgL)). A table comparing cohort sizes, differences in PFS and OS hazard ratios (HR) and significance (P) is shown (right). **b,** PFS and OS comparing IMiD treatment and t(IgH) status as in part a. P-values were determined by a Cox proportional hazards Wald’s test.

## Discussion

Here we present the first comprehensive catalogue of translocations in newly diagnosed multiple myeloma using whole genome sequencing on 826 tumor specimens with matched germline controls. Importantly, these results are placed in the context of clinical covariates and outcome, which allows for the identification of high-risk genetic alterations. This analysis identified 19 translocation hotspots present in ≥2% of the population and 8 recurrent translocations. As expected, the most common translocated region was IgH, which was primarily juxtaposed to known myeloma oncogenes *CCND1*, *WHSC1*, *MYC*, and *MAF.* Sequencing analysis provided high resolution of breakpoints and indicated that IgH translocations commonly occurred in the switch regions of the constant chains, implicating an error in class switch recombination in activated B cells, consistent with observations made over 20 years ago ^10^. However, this was not the case for IgH-MYC translocations, which were primarily located in extragenic regions further 3’ of the IgH locus, an observation made possible by the large number of samples analyzed here. This suggests that MYC translocations occur by a different mechanism and is consistent with the suggestion that MYC translocations are secondary events in myelomagenesis ^30^, albeit common secondary events. Additionally, t(11;14) breakpoints showed only marginal enrichment for class switch recombination regions, as several translocations also occurred in the VD and extragenic regions, similar to previous reports ^31^. This suggests that t(11;14) myelomas occur through at least two different mechanisms and may have distinct disease biology.

MYC translocations were present in 22.5% of newly-diagnosed patients and MYC amplification occurred in 15.6% of patients with half of those also containing a translocation, bringing the total number of newly diagnosed patients with MYC alterations to at least 30.6%. Both alterations resulted in commensurate upregulation of MYC as compared to other myelomas, but even those without MYC translocation or amplification, expressed MYC at a substantial level supporting the notion that MYC expression is a common feature of myeloma ^12^. The one exception to this are those myelomas with mutations in the MYC binding partner MAX, which express MYC at substantially lower levels ^32^. However, only MYC-IgL translocations experienced a poor prognosis, despite having similar baseline *MYC* expression as other MYC translocations. This pathology appeared to be intrinsic to the IgL locus as patients with IgL-translocated to non-MYC loci also experienced poor outcomes. This was further supported by the observation that t(IgL) myeloma patients had indistinguishable patient characteristics, clinical features, treatment regimens, and mutations as the entire cohort. This supports the notion that the IgL locus, when translocated to an oncogene, may be the driving feature of poor outcome.

Several studies have reported IgL translocations ^12,33–36^ but to our knowledge this is the first study in such a large cohort (N=826) of newly diagnosed myeloma and to report a frequency approaching 10%. This high frequency of IgL translocations was somewhat surprising given that 65% of myelomas express IgK, which was only translocated in 4.3% of patients. However, studies in murine B cells, 95% of which express IgK, found equivalent numbers of IgL and IgK translocations after acute activation with minimal time for oncogenic selection ^37^, suggesting that IgL is prone to translocation regardless of expression. Interestingly, the RNA-seq data here indicated that t(IgL) myeloma had the expected ratio of light chain expression with two-thirds of t(IgL) myeloma expressing IgK. This somewhat surprising result is consistent with the IgL enhancer being active even in IgK-expressing myeloma, which is supported by H3K27ac ChIP-seq data in IgK-expressing KMS11 and IgL-expressing MM.1S (**Fig. S9a,b**). However, 100% of IgK translocated myeloma expressed IgK (**Fig. S9c**). This observation can be explained by the fact that at least 75% of IgL-expressing B cells somatically delete the IgK constant region including the IgK 3’ enhancer ^38^, thus minimizing the potential for translocation and/or subsequent oncogenic propagation. Indeed, copy number data derived from the whole genome sequencing here clearly indicated that the majority of IgL-expressing myelomas had somatically deleted the IgK 3’ enhancer (**Fig. S9d**). Conversely, the IgL enhancer, which is one of the strongest enhancers in B cells, plasma cells, and myeloma, appears to be active regardless of whether or not a productive IgL is expressed as H3K27ac is found at IgL in IgK-expressing myeloma cell lines. Indeed, ‘super-enhancers’ are particularly prone to AID-induced genomic instability and translocation ^39^ and the IgL enhancer has been reported to be the strongest super-enhancer in myeloma ^40^. Subsequently, the IgL enhancer represents an unchecked and potent inducer of gene expression, that when transposed near an oncogene, robustly drives oncogenesis.

Although IgL translocations did not coincide with other small insertions/deletions, mutations, or patient clinical characteristics, t(IgL) was more common in hyperdiploid disease. The poor prognosis of t(IgL) is independent of hyperdiploidy and other major copy number alterations, which is exemplified by the additive risk when concurrent with amp(1q). This indicates that t(IgL) and amp(1q) function in independent pathways. Consequently, patients who harbor both of these high-risk genetic alterations experience an exceedingly poor outcome. Identification of t(IgL) myeloma will require a rapid and reliable diagnostic that can be used for appropriate risk-stratification. This is particularly important as t(IgL) is rarely identified clinically, whereas FISH is routinely performed to identify hyperdiploidy, which corresponds with better prognosis ^28^. Resultantly, t(IgL) represents an unrecognized high-risk marker prevalent in a subset of patients that are considered to be standard risk. Resultantly, the majority of t(IgL) are currently being misclassified as standard risk.

One indication of how t(IgL) may contribute to outcome is provided by the interaction of t(IgL) and treatment, which indicated that t(IgL) patients did not benefit from IMiDs as much as other translocated myelomas. The therapeutic efficacy of IMiDs may be mediated, in part, by their ability to inhibit expression from IKZF1 regulated enhancers, including those enhancers juxtaposed to oncogenes by translocation. While translocation of the immunoglobulin loci is a common mode of oncogene overexpression in myeloma, the IgL locus is unique in that, regardless of IgL expression, the IgL enhancers appear to be very active. This may render the IgL locus particularly insensitive to IMiD-based inhibition and potential explain why t(IgL) patients do not benefit from IMiDs. However, it will be important to further understand if and how IKZF1 regulates the IgL enhancers in the context of translocation. Recent work has shown the IgL locus to be one of the largest super-enhancers as measured by MED1 or BRD4 occupancy in MM.1S cells ^40^, suggesting several distinct transcriptional regulators may be mediating IgL-driven oncogene expression in t(IgL) myeloma. Thus, it will be important to understand the efficacy of emerging transcriptional therapeutics such as BET inhibitors ^41^ or degraders ^42^ that abolish BRD4 as well as other small molecule inhibitors that target the transcriptional machinery that drive oncogene expression. The data herein provide motivation for determining the efficacy of such transcriptional regulators in the context of t(IgL) myeloma, but also a rationale for the better understanding of cis-regulatory factors affecting transcription at translocated loci. This daunting task not only requires an understanding of the combinatorial effects of the trans-acting molecular machinery, but also a cartography of the chromatin structure, epigenetic mechanisms known to influence plasma cell fate ^43,44^, and enhancer function in the context of translocation breakpoints and micro-environmental ques. Such results will ultimately need to be placed in the context of multicellular organisms and bone marrow micro-environmental ques to help identify drug targets with high specificity for the transcriptional program and genetic architecture of myeloma.

## Methods

### CoMMpass data

Use of CoMMpass data (interim analysis 11) was approved by the data access use committee and downloaded from dbGaP (phs000748.v6.p4). Summarized data was provided by TGen.

### Whole genome sequencing

Long-insert whole genome library preparation and sequencing was performed by the Translational Genomics Institute (TGen) using 200-1,100 ng of DNA. DNA was sonicated (Covaris) to an average size of 900 bp. Libraries were prepared with either the TruSeq DNA Sample Prep (Illumina) or Hyper Prep (Kapa Biosystems) following the manufacturer’s instructions using AMPure beads (Invitrogen) for cleanup. Adapter ligated libraries were run on a 1.5% agarose gel and extracted by hand or with an Extractagel-extractor (USA Scientific). Size-selected libraries were purified with DNA Gel Extraction spin columns (Bio-Rad) prior to library amplification. Library size was assessed on a Bioanalyzer (Agilent) and quantified using the Qubit dsDNA HS assay (Invitrogen) and on a TapeStation (Agilent). Paired-end sequencing was performed using a HiSeq2000 or HiSeq2500 (Illumina) with either v3 or v4 chemistry and 86 bp reads. Raw sequencing data was extracted from BCL files using BCL2FASTQ v2.17.1 to extract FASTQ files.

### Whole genome sequencing analysis

Sequencing FASTQ files were mapped to a modified version of the human genome GRCh37 using BWA (v0.7.8) ^45^. The specific reference genome was phase 2 of the 1,000 Genomes project ^46^ plus contigs from ribosomal DNA and cancer-related viruses (U13369.1, FR872717.1, AF092932.1, NC_001526.2, NC_001357.1, NC_003977.1, NC_004102.1, NC_009823.1, NC_009824.1, NC_009825.1, NC_009826.1, NC_009334.1, NC_006273.2, NC_011800.1, NC_001806.1) as well as sequence from the External RNA Controls Consortium (ERCC) ^47^. Aligned BAM files were sorted with SAMtools (v0.1.19) ^48^, recalibrated with GATK (v3.1.1) ^49^, and putative PCR duplicates were marked with Picard (v1.111). Tumor-specific indel realignment was performed relative to normal samples collected at the same time from the same patient using GATK (v3.1.1).

### Determination of structural variants

Structural variants were determined using DELLY (v0.7.6) ^22^ with the following filter module options: altFrac = 0.1, ratioGeno = 0.75, coverage = 5, controlContamination = 0, minSize = 500, maxSize = 500000000. DELLY BCF files were converted to VCF files using BCFTOOLS and VCF files were parsed, formatted, and annotated using custom R scripts available upon request. DELLY translocations were subject to further quality control that included homology searches of 1 kb on either side of the translocation break point between the two transposed regions where translocations with homology of 80% or more in any 100 bp window were removed as false positives. Translocations were compared to the ENCODE ^50^ 20bp mappability tracks and those translocations that had an average mappability of less than 20% across 1 kb on either side of the translocation breakpoint were also removed. Finally, translocations were visually inspected and compared to the mapped reads in BAM files resulting in elimination of translocations in certain regions with sequencing anomalies (**Table S6**).

### Identification of translocation hotspots

Commonly translocated regions were identified using a 1 Mb window incremented 500 kb across each chromosome. Contiguous 1 Mb regions translocated at a frequency of 1% or more were stitched together. Common translocations were identified by calculating the frequency of any given translocation type within 1 Mb of the two chromosomal breakpoints.

### Translocation visualization

R / Bioconductor (v3.4.3) ^51^ was used to generate circos plots with the RCircos package (v1.2.0) ^52^. Heatmaps of translocation occurrence by sample were generated with the ‘image’ function in a method similar to the ‘heatmap’ function of the stats package. Plots of translocations across genomic regions were made as previously described ^43^ and translocations were represented by a region covering 1 kb of each side of the breakpoint or estimated breakpoint.

### RNA-seq

RNA-seq libraries were constructed with the TruSeq RNA Library Prep Kit v2 (Illumina) by TGen, which yields unstranded mRNA libraries. 150-2,000 ng of RNA, which had an RNA integrity number (Agilent Bioanalyzer) of 8 or higher was used for starting material. RNA libraries were amplified for 8-10 cycles and then sequenced on an Illuina HiSeq2000 or HiSeq2500 using v3 or v4 chemistry and 82bp paired-end reads.

### RNA-seq analysis

RNA-seq reads were aligned to the same GRCh37 genome as whole genome sequencing data using STAR (v2.3.1) ^53^ with the GRCh37.74 gtf file to guide splice junction identification. Coverage was determined with HTSeq (v0.6.0) ^54^ and FPKM normalization was done using R / Bioconductor without the use of IMGT-defined immunoglobulin genes, which were analyzed separately. IgH constant chain expression was determined by the most highly expressed constant region so long as that was ≥5,000 FPKM. Likewise, light chain expression was determined as the highest expression (measured by FPKM) of the cumulative variable, joining, and constant regions for IgK and IgL. Expression of individual genes were plotted with R ‘boxplot’ function with outliers plotted using the beeswarm package (v0.2.3). Ad-hoc differences in gene expression for translocated genes was determined with the Mann-Whitney U-test for pairwise comparisons or analysis of variance with Tukey’s post-hoc correction for multiple comparisons.

Gene expression subtypes were determined using the ‘ConcensusClusteringPlus’ (v1.42.0) ^55^ in R/Bioconductor using log_2_(FPKM+1) transformed data that was gene normalized for 1,000 iterations using a hierarchical clustering algorithm and a Pearson distance metric performed on 2 to 20 consensus groups. Increasing the consensus groups beyond 7 explained only minimal variation, thus 7 was chosen. Genes uniquely expressed in each consensus group were determined by comparing each group to all other samples using edgeR ^56^ with an FDR ≤0.01 as were genes differentially expressed between t(IgL) myeloma and other myelomas. t(IgL) differentially expressed genes were further confirmed by restricting the analysis to only samples with hyperdiploidy and by performing analysis with a co-variate for gene expression subtype determined by consensus clustering. Gene set enrichment analysis (GSEA) ^25^ was performed using a pre-ranked list determined by the −log_10_(P-value) × sign(fold-change) against all curated gene lists in MSigDB (v6.1) and for those from Zhan *et al.* ^24^ to match gene expression subtypes to those previously identified.

### Exome-sequencing

Exome sequencing was performed by TGen on DNA from CD138+ myeloma cells and peripheral blood references samples for all individuals. DNA was fragmented with a Covaris and library preparation was performed with either TruSeq exome kit (Illumina) or with the HyperPrep kit (Kapa) in combination with the Sureselect V5+UTR exome capture kit (Agilent). Data were mapped to the same reference genome (GRCh37, plus contigs defined above) using BWA (v0.7.8) ^45^. As with the whole genome long-insert sequencing, aligned BAM files were sorted with SAMtools (v0.1.19) ^48^, recalibrated with GATK (v3.1.1) ^49^, and putative PCR duplicates were marked with Picard (v1.111).

### Mutational analysis

Somatic SNV and INDELS were based on calls from Seurat (v2.6) ^57^, Strelka (v1.013) ^58^, and MuTect (v1.1.4) ^59^. Final mutations were determined as those that were called by at least 2 of the 3 aforementioned tools. Mutations were annotated to GRCh37.74 protein coding genes and immunoglobulin genes were excluded from analysis due to somatic hypermutation. Synonymous and non-synonymous mutations were determined using the ‘locateVariants’ and ‘predictCoding’ functions of the ‘VariantAnnotation’ (v1.24.5) package ^60^. Mutational differences between t(IgL) and non-t(IgL) myelomas were assessed using Fisher’s exact test with an FDR correction for all mutations present at frequency of ≥4% of the population.

### Copy number alteration analysis

Copy number alterations (CNAs) were determined separately for exome sequencing and whole genome long-insert sequencing using the TGen tool tCoNut (https://github.com/tgen/tCoNuT). Exome-sequencing derived CNAs were used to define large cytogenetic abnormalities including hyperdiploidy, del(1p), amp(1q), del(13q), and del(17p). CNA gains and losses were defined as log_2_ CNA ratio of myeloma to normal of ≥0.2 and ≤0.2, respectively. The following regions were used to define common myeloma CNAs based on the average CNA segmentation call for the region.

| Name | Region |
| --- | --- |
| del(1p) | 1p22 |
| amp(1q) | 1q21 |
| del(13) | 13q14 |
| del(17) | 17p13 |
| Hyperdiploidy* | 3p22, 3q21, 5p15, 5q31, 7p14, 7q22, 9p13, 9q33, 11p15, 11q23, 15q15, 15q26, 19p13, 19q13, 21q21, 21q22 |

\*Hyperdiploidy required concordant gains of both areas on a given chromosome for 4 chromosomes.

CNA analysis of global chromosome changes (*i.e.* Fig. 6A) was done by calculating the average CNA log_2_ ratio for 100 kb bins across the genome and clustering was performed using ConsensusClusterPlus ^55^. Intergenic CNAs, including those at the MYC, IgK, and IgL loci, were derived from whole genome long-insert sequencing data and binned into 100 bp increments.

### Survival analysis

Survival analysis was conducted using in R using the ‘survival’ (v2.41-3) package. Differences in progression-free and overall survival were determined using a cox proportional hazards regression fit to either a continuous (e.g. the number of deletions, duplications, inversions, or translocations) or discrete (e.g. t(IgL) versus other) variable. P-values were calculated using a Wald’s test. When more than two discrete variables existed a P-value of differences between all groups was first calculated followed by pairwise comparisons and FDR correction. Hazard ratios associated with translocations, mutations, and other clinical variates were also calculated using a cox proportional hazards regression and 95% confidence intervals are shown. Bivariate analysis was performed for the most common mutations in combination with t(IgL) and multivariate analysis was conducted with clinically relevant parameters and t(IgL) as well as common CNAs and t(IgL). Bootstrapping of outcome was performed using 1,000 permutations and comparing the PFS and OS hazard ratios as compared to the actual hazard ratio. The comparison of IMiD survival benefit in t(IgL) versus t(IgH) sampled t(IgH) patients according to the number of t(IgL) patients.

### ChIP-seq analysis

H3K27ac ChIP-seq data generated as part of the ENCODE project ^50^ for the myeloma cell lines KMS11 and MM.1S or by Lin et al. ^29^ for MM.1S were downloaded from the short read archive and mapped to the same GRCh37 genome used above for CoMMpass data using bowtie 2 (v2.2.6) ^61^. Mapped SAM files were converted to BAM files and putative PCR duplicates were marked using SAMtools (v1.7) ^48^. H3K27ac and IKZF1 enriched regions were determined using MACS2 (v2.1.0.20151222) ^62^ using default parameters and a q-value of 0.01. Fragment size was estimated using the R package ‘chipseq’ (v1.28.0) and reads were extended to the estimated fragment size for visualization using the R package ‘rtracklayer’ (v1.38.3) ^63^. Super-enhancer analysis was performed using custom R code in a manner analogous to that done previously ^40^. Briefly, this involved stitching together IKZF1-enriched regions that were within 15 kb of each other for enriched regions that did not overlap a 2.5 kb proximal to a GRCh37.74 defined promoter. Regions were then ranked by IKZF1 occupancy measured as reads per million (RPM) and regions that were past the inflection point were considered super-enhancers.

### Data availability

CoMMpass data is deposited in dbGaP (phs000748.v6.p4) and summarized data can be accessed at https://research.themmrf.org/.

### Code availability

All code is available upon request.

## Acknowledgments

We wish to thank the patients, researchers and clinicians involved in the Multiple Myeloma Research Foundation CoMMpass trial. We thank Dr. Madhav V. Dhodapkar for helpful critique. This work was supported by a Precision Medicine award from the MMRF to BGB, SL, PMV, and LHB. BGB is supported by Postdoctoral Fellowship PF-17-109-1-TBG from the American Cancer Society.

## Author Contributions

BGB performed bioinformatic analysis, interpreted the data, and wrote the paper. PN and NJB performed IKZF1 ChIP-seq and help interpret the data. AKN, JLK, and VAG interpreted the data. DA provided administration for the CoMMpass trial and interpreted the data. JJK oversaw genomic and bioinformatic analyses, SL was the clinical PI of the CoMMpass study, interpreted data, and help write the manuscript. PMV and LHB interpreted data and wrote the manuscript.

## Competing Interests

The authors declare no competing interests.

